# Ascertainment bias in the genomic test of positive selection on regulatory sequences

**DOI:** 10.1101/2023.08.20.554030

**Authors:** Daohan Jiang, Jianzhi Zhang

**Affiliations:** Department of Ecology and Evolutionary Biology, University of Michigan, Ann Arbor, Michigan 48109, USA

**Author notes:** Correspondence to Jianzhi Zhang, Department of Ecology and Evolutionary Biology, University of Michigan, 4018 Biological Sciences Building, 1105 North University Avenue, Ann Arbor, MI 48109, USA. Department of Quantitative and Computational Biology, University of Southern California, Los Angeles, California 90089, USA.

**Keywords:** transcription factor binding sites, human, adaptation, ChIP-seq, regulatory evolution

## Abstract

Evolution of gene expression mediated by *cis*-regulatory changes is thought to be an important contributor to organismal adaptation, but identifying adaptive *cis*-regulatory changes is challenging due to the difficulty in knowing the expectation under no positive selection. A new approach for detecting positive selection on transcription factor binding sites (TFBSs) was recently developed, thanks to the application of machine learning in predicting transcription factor (TF) binding affinities of DNA sequences. Given a TFBS sequence from a focal species and the corresponding inferred ancestral sequence that differs from the former at *n* sites, one can predict the TF binding affinities of many *n*-step mutational neighbors of the ancestral sequence and obtain a null distribution of the derived binding affinity, which allows testing whether the binding affinity of the real derived sequence deviates significantly from the null distribution. Applying this test genomically to all experimentally identified binding sites of three TFs in humans, a recent study reported positive selection for elevated binding affinities of TFBSs. Here we show that this genomic test suffers from an ascertainment bias because, even in the absence of positive selection for strengthened binding, the binding affinities of known human TFBSs are more likely to have increased than decreased in evolution. We demonstrate by computer simulation that this bias inflates the false positive rate of the selection test. We propose several methods to mitigate the ascertainment bias and show that almost all previously reported positive selection signals disappear when these methods are applied.

## INTRODUCTION

Organism-level phenotypic changes in adaptive evolution are believed to be often caused by gene expression alterations brought by changes in *cis*-regulatory sequences (King and Wilson 1975; Wray 2007; Carroll 2008; Jones, et al. 2012; Signor and Nuzhdin 2018). However, it is challenging to test positive selection on *cis*-regulatory sequences because the neutral expectation is difficult to know, unlike the test of positive selection on protein-coding sequences where the neutral expectation is usually assumed to be reflected by synonymous substitutions (Li, et al. 1985; Nei and Gojobori 1986; McDonald and Kreitman 1991; Nei and Kumar 2000). While one can test positive selection on the *cis*-regulatory sequence of a gene by comparing it with synonymous sites in the gene (Andolfatto 2005), the comparison would rely on the assumption that the regions being compared have equal mutation rates. The non-neutrality of many synonymous mutations (Lind, et al. 2010; Lawrie, et al. 2013; Sharon, et al. 2018; She and Jarosz 2018; Shen, et al. 2022) further complicates the test. Comparing the *cis*-regulatory sequence of a gene with the intron sequences of the gene is another choice (Haygood, et al. 2007), but it similarly depends on the assumption that the regions being compared have equal mutation rates and that introns evolve neutrally. Additionally, because the number of functional sites in a *cis*-regulatory sequence is typically small, the above comparisons are generally statistically underpowered.

Because functional changes of *cis*-regulatory sequences typically occur via altering their bindings to *trans*-regulatory factors such as transcription factors (TFs), selection on *cis*-regulatory sequences is in a large part mediated by selection on binding affinities (Berg, et al. 2004; Moses 2009). In theory, one can compare an observed evolutionary change in binding affinity with a null distribution in the absence of selection to test whether the *cis*-regulatory sequence has been positively selected. Such a test is possible only if the binding affinities of numerous potential mutant sequences, which would be labor-intensive to quantify experimentally, are known. This problem was recently solved by using machine learning to predict TF binding affinities (Liu and Robinson-Rechavi 2020). Specifically, based on previously established techniques (Ghandi, et al. 2014; Lee, et al. 2015; Ghandi, et al. 2016), Liu and Robinson-Rechavi trained a gapped *k*-mer support vector machine (gkm-SVM) using TF binding sites (TFBSs) experimentally identified through chromatin immunoprecipitation followed by sequencing (ChIP-seq). This trained program is then used to calculate SVM weights (i.e., contribution to the overall binding affinity) of all possible 10-mers, based on which a binding affinity score (SVM score) can be calculated for an arbitrary sequence by summing up SVM weights of all 10-mers that it contains.

With the above tool, one can predict the affinity scores of many mutational neighbors of an ancestral regulatory sequence to obtain a null distribution of binding affinity changes and compare the observed evolutionary change (deltaSVM) with this null distribution. If the observed deltaSVM is in the right 5% or 1% tail of the null distribution, one could conclude that positive selection for a higher binding affinity to the TF of interest acted in the evolution of the regulatory sequence (Prabhakar, et al. 2006; Gittelman, et al. 2015). For example, based on this method, Liu and Robinson-Rechavi (2020) reported positive selection for elevated binding affinities in human evolution since the human-chimpanzee split for a few percent of the binding sites of each of three TFs examined: CEBPA, HNF4A, and CTCF.

In such tests of positive selection, the TFBSs considered are typically acquired from ChIP-seq peaks (Park 2009) identified in a focal species such as the human in the above study. This means that only TFBSs with relatively high binding affinities are included in the analysis, which could cause an ascertainment bias because binding affinities are more likely to have increased for TFBSs with relatively high affinities than for TFBSs with relatively low affinities even in the absence of positive selection for higher affinities (**Fig. 1**). As a result, one may miscall chance increases of binding affinities as signals of positive selection, raising the false positive rate in the test of positive selection. An analogy of this problem is to test whether students generally perform better in the final exam than in the midterm exam while considering only those who perform well in the final exam.

**Figure 1.**
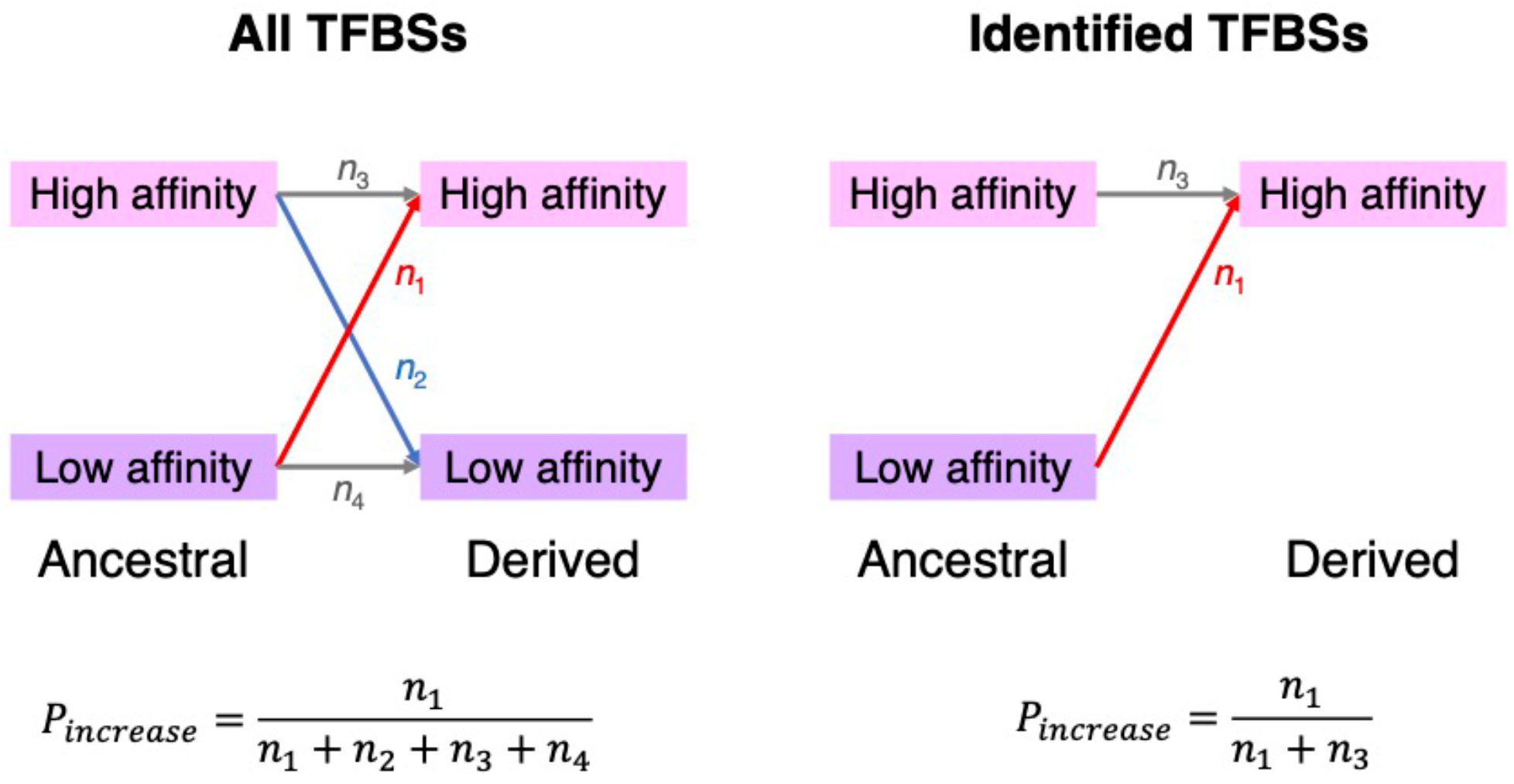
Schematic illustration of the ascertainment bias in the selection test of Liu and Robinson-Rechavi (2020). The left panel represents the scenario where all potential TFBSs in an ancestor are included in the tests so there is no ascertainment bias, whereas the right panel represents the scenario in the actual test of Liu and Robinson-Rechavi where there is an ascertainment bias. Here, *n*_1_, *n*_2_, *n*_3_, and *n*_4_ denote the number of potential TFBSs that underwent each type of binding affinity change, respectively. *P*_increase_ denotes the observed proportion of TFBSs with increased binding affinities and is unbiasedly estimated in the left panel. In the right panel, however, *P*_increase_ is overestimated because TFBSs with low derived affinities (in the focal species) are not included.

In this study, we use computer simulation of neutral evolution to demonstrate the influence of the ascertainment bias on the test of positive selection on TFBSs. We propose three methods to mitigate this impact and show that almost all signals of positive selection previously reported for the binding sites of the three human TFs (Liu and Robinson-Rechavi 2020) disappear upon ascertainment bias mitigations.

## RESULTS

### Interpretation of *P*-values in the selection test

For each TFBS subject to Liu and Robinson-Rechavi’s one-tailed test of positive selection for elevated binding affinity, the *P*-value is the probability that deltaSVM under neutrality (denoted Δ for short) is greater than the observed value (Δ_obs_), given the ancestral sequence and the number of nucleotide substitutions separating the ancestral and derived sequences. That is, *P* = *Prob*(Δ > Δ_obs_). However, for a TFBS from a real dataset, the probability that it shows Δ >_obs_ under neutrality is the conditional probability 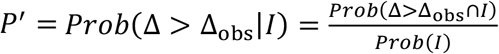, where *I* denotes the event that the TFBS is included in the dataset. Because TFBSs with higher Δ_obs_ should have higher *Prob*(*I*) but lower *Prob*(Δ > Δ_obs_), events *I* and Δ > Δ_obs_ are not mutually independent but tend to avoid each other. That is, *Prob*(Δ > Δ_obs_ ∩ *I*) < *Prob*(Δ > Δ_obs_)*Prob*(*I*). Thus, 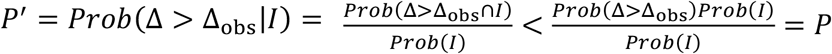. This inequation shows that *P* will be underestimated if *P’* is interpreted as *P*. Under neutrality, if the probability for *P* < 0.01 is 0.01, the probability for *P’* < 0.01 must be greater than 0.01, inflating the false positive rate in the test of positive selection. If the test is unbiased, the action of positive selection can be inferred when the proportion of TFBSs showing *P* < 0.01 is greater than 1%. However, this inference is no longer valid if *P’* instead of *P* is calculated because the fraction of TFBSs showing *P’* < 0.01 under neutrality is expected to exceed 0.01. Therefore, the ascertainment bias is a problem both in assessing positive selection on an individual TFBS and on a dataset containing many TFBSs.

### False positive rates under various levels of the ascertainment bias

We performed a computer simulation to evaluate the quantitative impact of the ascertainment bias on the false positive rate. We first generated 50,000 10-nucleotide random sequences as ancestral TFBS sequences. In each sequence, we then introduced two random substitutions at two randomly picked sites to generate the corresponding derived sequence. We treated these sequences as potential binding sites of human CEBPA studied by Liu and Robinson-Rechavi. To mimic the ascertainment bias, we removed 10%, 20%, …, and 90% of the simulated TFBSs with the lowest derived SVM scores, obtaining nine datasets with increasing ascertainment biases. We then performed a one-tailed test of positive selection for an elevated CEBPA binding affinity for each TFBS. Following Liu and Robinson-Rechavi, we called a case statistically significant when the right-tail probability is below 0.01 (see Materials and Methods). Because all substitutions introduced are random, all cases of positive selection identified are false positives. Indeed, each of the nine datasets with ascertainment biases showed a significantly higher false positive rate than that of the original dataset without the ascertainment bias (*P* < 0.05, two-sample *Z*-test for proportions), and the false positive rate increased with the severity of the ascertainment bias (**Table 1**). For the most extreme level of ascertainment bias considered, the false positive rate is 11 times the expected level. We repeated the above analysis by treating the sequences as potential binding sites of two other human TFs studied by Liu and Robinson-Rechavi (HNF4A and CTCF) and obtained highly similar results (**Tables S1-S2**).

**Table 1.** All simulated TFBSs and subsets with various fractions of the TFBSs with the highest derived SVM scores for human CEBPA. *P*-value in the last column is from a test of the null hypothesis that the false positive rate in a subset equals that in the complete dataset (two-sample *Z*-test for proportions).

| Subset | No. of TFBSs | No. of TFBSs with positive deltaSVMs | No. of false positives | *P*-value |
| --- | --- | --- | --- | --- |
| Complete dataset | 50,000 | 24,908 (49.82%) | 553 (1.106%) | N.A. |
| Top 90% | 45,000 | 24,380 (54.18%) | 553 (1.230%) | $4.2 \times 10^{-2}$ |
| Top 80% | 40,000 | 23,281 (58.20%) | 553 (1.383%) | $1.0 \times 10^{-4}$ |
| Top 70% | 35,000 | 21,810 (62.31%) | 553 (1.580%) | $1.2 \times 10^{-9}$ |
| Top 60% | 30,000 | 20,003 (66.68%) | 553 (1.843%) | $3.5 \times 10^{-18}$ |
| Top 50% | 25,000 | 17,748 (70.99%) | 553 (2.212%) | $1.7 \times 10^{-32}$ |
| Top 40% | 20,000 | 15,085 (75.43%) | 553 (2.765%) | $5.3 \times 10^{-57}$ |
| Top 30% | 15,000 | 12,012 (80.08%) | 553 (3.687%) | $6.9 \times 10^{-102}$ |
| Top 20% | 10,000 | 8,464 (84.64%) | 553 (5.530%) | $8.2 \times 10^{-198}$ |
| Top 10% | 5,000 | 4,448 (88.96%) | 552 (11.040%) | $< 10^{-300}$ |

### Methods for mitigating the ascertainment bias

With the ascertainment bias, the expected false positive rate under neutrality is unknown for a given dataset. Hence, one cannot directly use the proportion of TFBSs in a dataset that show significant positive selection signals to infer whether positive selection has occurred.

Rather, one should also consider the TFBSs that are not included in the data due to the ascertainment bias. Our simulation showed that it is virtually impossible for any of these missing TFBSs to show significant positive selection signals (**Table 1, Tables S1-S2**). Hence, the missing TFBSs need only be added to the denominator when one computes the proportion of TFBSs with significant positive selection signals. That is, the corrected proportion of TFBSs with significant positive selection signals equals the number of TFBSs with such signals divided by the sum of considered and missing TFBSs. Alternatively, one can infer the occurrence of positive selection on an individual TFBS by using adjusted *P*-values after correcting for multiple testing, where the number of tests should be the sum of the considered and missing TFBSs.

Below we present three different methods for estimating the number of missing TFBSs.

The first method regards twice the number of TFBSs with positive deltaSVMs as the total number of tests, because the number of TFBSs with negative deltaSVMs should be similar to that with positive deltaSVMs in the absence of ascertainment bias. Note that this method likely still undercounts the total number of tests, because it cannot recover TFBSs with low binding affinities in both the ancestral and focal taxa (i.e., *n*_4_ in **Fig. 1**).

The second method first identifies the relationship between the fraction of TFBSs removed and the fraction of TFBSs with positive deltaSVMs in our simulated data (**Table S3, Fig. S1**) and then uses this relationship to infer the fraction of missing TFBSs and the total number of tests for a real dataset. Note that this method may not accurately recover the number of missing TFBSs for the following reason. Biased subsets of our simulated dataset were obtained by removing TFBSs with the lowest derived SVM scores. In reality, however, loss of TFBSs with low derived affinities likely follows a different yet unknown model, because TFBSs with low binding affinities can sometimes be detected, albeit with a relatively low probability.

Consequently, this correction method likely underestimates missing TFBSs and therefore under-corrects the ascertainment bias.

The third method regards the number of potential binding sites of a given TF in the genome that have at least two nucleotide differences (between the ancestral and derived sequences) as the number of total tests. If the binding sites for the TF is on average *l* nucleotides long, there are (*L*-*l*+1)*P*(*n* ≥ 2) ≈ *LP*(*n* ≥ 2) total tests, where *L* >> *l* is the total number of nucleotides in the genome and *P*(*n* ≥ 2) is the fraction of TFBSs that have at least two nucleotide differences between the ancestral and derived sequences (see Materials and Methods). In the above calculation, potential binding sites are overlapping. However, to be more conservative in computing the total number of tests, we may require the potential binding sites to be nonoverlapping, which yields an estimate of *LP*(*n* ≥ 2)/*l* potential binding sites in the genome.

### Most positive selection signals disappear upon ascertainment bias mitigations

As mentioned, Liu and Robinson-Rechavi (2020) reported the detection of positive selection on the binding sites of each of three human TFs investigated, because >1% of the TFBSs fall in the right 1% tail of the null distribution of deltaSVM. The TFBSs examined were all identified from the human ChIP-seq data, so the test suffered from ascertainment biases.

Below we use the three methods proposed in the preceding section to mitigate the ascertainment bias and re-evaluate the evidence for positive selection.

In the case of CEBPA, the original study reported that, of 5,807 TFBSs tested, there were 436 (or 7.51%) that showed positive selection signals (referred to as positive binding sites, or PBSs for short following terminology used by Liu and Robinson-Rechavi) at the 1% significance level. Based on the first method of correction, the total number of tests should be twice the number of TFBSs with positive deltaSVMs, or 2×3618 = 7236. Hence, the proportion of PBSs declines to 436/7236 = 6.03% (**Table 2**). Under the second method of correction, the total number of tests is estimated to be 8200 (**Table S3**), leading to 436/8200 = 5.32% of PBSs (**Table 2**). Considering the third method of correction and nonoverlapping TFBSs, we estimated that that there are 1,088,177 potential TFBSs, meaning that 436/1088177 = 0.0401% of TFBSs are PBSs (**Table 2**). In the case of HNF4A, the fraction of PBSs is 4.81% before correction for ascertainment bias and reduces to 4.11%, 3.83%, and 0.0324% when methods 1, 2, and 3 are used, respectively (**Table 2**). In the case of CTCF, the fraction reduces from 3.52% before correction to 3.20%, 3.12%, and 0.0375% after the three correction methods are used (**Table 2**). After the third correction, the fraction of PBSs detected is below the neutral expectation of 1% for each of the three TFs, suggesting that positive selection is not needed to explain the evolution of the binding sites of the three TFs.

**Table 2.** Number and fraction of TFBSs with signals of positive selection (i.e., positive binding sites or PBSs) before correction for multiple testing. Denominator used to calculate the fraction is either the observed number of TFBSs or the inferred number of TFBSs including those that are missing due to ascertainment biases.

|  | **CEBPA binding sites** | **HNF4A binding sites** | **CTCF binding sites** |
| --- | --- | --- | --- |
| Observed number | 5,807 | 12,734 | 80,074 |
| Number with positive deltaSVMs | 3,618 | 7,438 | 44,117 |
| Fraction with positive deltaSVMs | 62.15% | 58.41% | 55.10% |
| Number of PBSs | 436 | 612 | 2821 |
| Fraction of PBSs (no correction*) | 7.51% | 4.81% | 3.52% |
| Fraction of PBSs (method 1*) | 6.03% | 4.11% | 3.20% |
| Fraction of PBSs (method 2*) | 5.32% | 3.83% | 3.12% |
| Fraction of PBSs (method 3*) | 0.0401% | 0.0324% | 0.0375% |

\*The total number of TFBSs considered is the observed number of TFBSs in the dataset (no correction), twice the number of observed TFBSs with positive deltaSVMs (method 1), inferred from a linear regression model (method 2), or an estimate of the number of potential binding sites in the genome (method 3), respectively.

We similarly evaluated positive selection on individual TFBSs by estimating adjusted *P*-values. If we use the cutoff of adjusted *P*-value = 1%, there is no PBS when the ascertainment bias is uncorrected or corrected by any of the three methods (**Table 3**). If we use the cutoff of adjusted *P*-value = 5%, 109 CEBPA binding sites show significant positive selection signals when the ascertainment bias is uncorrected. This number reduces to 80, 58, and 0, respectively, under the three methods of correction (**Table 3**). Considering both the results from the proportion of TFBSs with significant positive selection signals and the individual TFBSs with such signals, we conclude that few if any human CEBPA binding sites evolved under positive selection. Binding sites of the other two human TFs show even weaker signals of positive selection. No HNF4A binding sites have adjusted *P* < 0.05 regardless of the method used to correct the ascertainment bias (**Table 3**). A total of 161 CTCF binding sites have adjusted *P* < 0.05 when method 1 is used to correct the ascertainment bias, but this number reduces to 0 when method 2 or 3 is applied (**Table 3**).

**Table 3.**
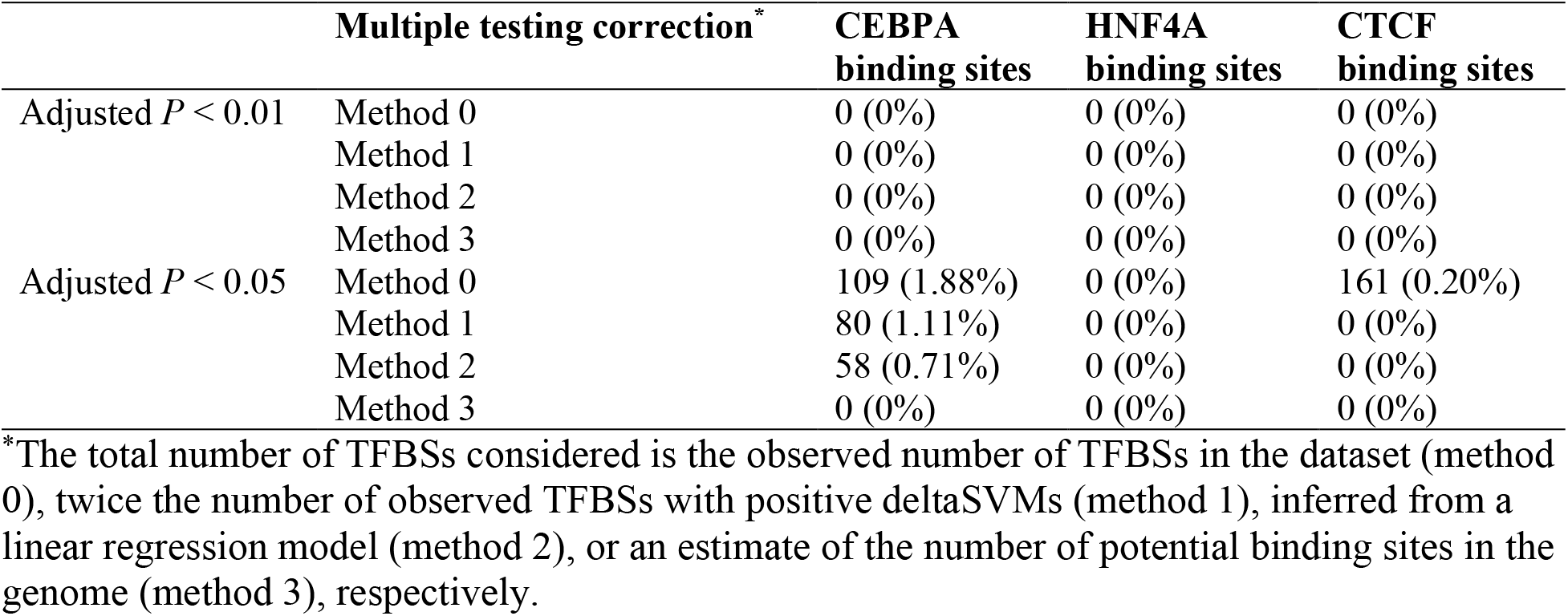
Number and fraction of positively selected TFBSs (PBSs) after correction for multiple testing. When the number of tests is corrected for, the fraction is calculated using the inferred number of TFBSs including those missing due to ascertainment biases.

## DISCUSSION

We showed that a previously proposed genomic test for positive selection on TFBSs for increased binding affinity suffers from inflated false positive errors due to ascertainment biases. Because such biases in the test are inevitable in real data, without an appropriate correction, the selection test would be unable to identify positively selected TFBSs accurately. After applying various corrections, we found no evidence for positive selection on the binding sites of human xHNF4A and CTCF and no to weak signals of positive selection on the binding sites of human CEBPA.

We considered three different methods to recover the missing TFBSs caused by the ascertainment bias. Methods 1 and 2 likely provide insufficient corrections for the following reasons. While method 1 probably recovers TFBSs that have detectable binding affinities in the ancestral node but not in the focal species (i.e., *n*_2_ in **Fig. 1**), it does not recover TFBSs that lack detectable binding in both species (i.e., *n*_4_ in **Fig. 1**). Method 2 utilizes the linear relationship observed in the simulated data between the fraction of TFBSs removed and the proportion of remaining TFBSs with positive deltaSVMs. However, when introducing the ascertainment bias in the simulation, we simply removed TFBSs with the lowest binding affinities. Given the target fraction of TFBSs with positive deltaSVMs, this approach minimizes the removed fraction of TFBSs. Because experimental identifications of TFBSs are likely subject to some noise, the actual missing TFBSs may not have the lowest binding affinities. Consequently, method 2 likely underestimates the number of missing TFBSs. This said, if there are already no significant positive selection signals after one applies methods 1 and 2, more stringent corrections would not be needed. If there are positive selection signals, however, they should be further tested by applying more rigorous corrections or be treated as candidates for further tests such as an experimental verification of fitness effects or population genetic test of selective sweeps.

The third correction method (method 3) considers the number of potential TFBSs in the whole genome. In theory, this method provides the most complete correction. However, estimating the total number of potential TFBSs (i.e., number of sequence segments that can potentially become TFBSs in evolution) can be challenging. In this study, we assumed that all genomic regions can potentially become a TFBS. This assumption is reasonable at least qualitatively, because “leaky expression” caused by fortuitous TF binding is indeed widespread, as evidenced by the observation that over three quarters of the human genome is transcribed and that about one half of 120-nucleotide random sequences can drive gene expression in yeast (Xu, et al. 2023). However, if different genomic regions have different potentials to evolve into a TFBS in the focal lineage, method 3 could overcorrect the ascertainment bias when regions with low potentials are counted. This said, if such heterogeneities can be modeled, method 3 should provide an appropriate correction of the ascertainment bias.

It should be noted that, because the *P*-value of the selection test is calculated based on a finite number of (simulated) mutants, the test inevitably loses power when multiple testing is corrected. For instance, Liu and Robinson-Rechavi (2020) considered 10^4^ mutants per test, so the smallest possible *P*-value was 10^−4^, yet the number of TFBSs in the dataset can be very large (e.g., on the order of 10^4^ for HNF4A and CTCF; **Table 2**).

Our re-analysis of the binding sites of three human TFs showed no signal of positive selection on the TFBSs of HNF4A and CTCF and no to weak signals of positive selection on the TFBSs of CEBPA (**Table 3**). The latter finding appears consistent with the validation tests Liu and Robinson-Rechavi performed on CEBPA binding sites. One of their validations, inspired by the McDonald-Kreitman test (McDonald and Kreitman 1991), compares the ratio of the number of substitutions to the number of polymorphisms between PBSs and non-PBSs. This test alone, however, is not a sufficient means of validation, because many non-PBSs are likely subject to negative selection because of stabilizing selection on their binding affinities, and negative selection can reduce the divergence-to-polymorphism ratio (Eyre-Walker 2002). Liu and Robinson-Rechavi further showed that the target genes of PBSs tend to have lowered expression variance across human populations, suggesting that the binding affinities of non-PBSs are not generally under stronger stabilizing selection than PBSs. The two validation tests, together with our reanalysis, suggest the possibility of positive selection on some CEBPA binding sites in the human lineage.

In addition to human TFBSs, Liu and Robinson-Rechavi also analyzed TFBSs in mice and fruit flies. These two datasets do not suffer from the ascertainment bias as severely as the human datasets suffer, because TFBSs detected in focal and non-focal species (or lineages) are all included. However, these datasets are not free of the ascertainment bias because only TFBSs with sufficiently strong binding in at least one species can be included, whereas proto-TFBSs that have low binding in all species considered (and presumably in their most recent common ancestor too) cannot. That is, the low-to-low class (*n*_4_) in **Fig. 1** are still missing, and the number of TFBSs in the dataset is still smaller than the size of the ideal “complete” dataset. As a result, selection tests have inflated false positive errors. For example, the fraction of TFBSs with increased binding should be 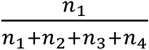 but is instead computed by 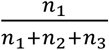 (see **Fig. 1**).

Because *n*_4_ is presumably the largest among the four *n* values, the above two fractions are very different. This problem can be mitigated to some extent by broader phylogenetic sampling, though ancestral sequence reconstruction would be subject to more errors as more divergent species are included.

We conclude that the previously proposed genomic test is unable to rigorously estimate the prevalence of positive selection on TFBSs due to an ascertainment bias. There are multiple ways to mitigate the bias, but it remains challenging to adequately correct but not overcorrect the bias. We suggest that it is necessary to combine multiple methods to verify the signal of positive selection on TFBSs, including broader phylogenetic sampling, correction for ascertainment bias, correction for multiple testing, and other means to test selection, to robustly estimate the prevalence of positive selection on TFBSs and identify individual TFBSs where nucleotide substitutions have been driven by positive selection.

Finally, it is worth pointing out that the type of ascertainment bias encountered here is not unique to the test of positive selection on TFBSs, especially at the post-genomic era.

Evolutionary analyses of subsets of genomic data that satisfy certain criteria are potentially subject to ascertainment biases when the criteria are correlated with the factors being investigated. A prime example is the test of Ohno’s hypothesis of X-chromosome dosage compensation in mammals (i.e., doubling of the expressions of X-linked genes to compensate the degeneration of their Y-linked counterparts) (Ohno 1967), where considering only genes with expression levels higher than a cutoff produced misleading results (He, et al. 2011; Lin, et al. 2012). Caution should be exerted in designing such tests.

## MATERIALS AND METHODS

### Analysis of simulated TFBSs

To generate a dataset that represents potential TFBSs, we randomly generated 50,000 distinct 10-nucleotide sequences with equal probabilities of the four nucleotides at each site. In each sequence, we randomly picked two sites and make a random change at each site such that the derived sequence differs from the original, ancestral sequence by two substitutions (with no multiple hits and no mutational bias).

Following Liu and Robinson-Rechavi (2020), for each pair of ancestral and derived TFBSs, we performed a one-tailed test of positive selection for an increased binding affinity of the derived TFBS. We separately considered three human TFs—CEBPA, HNF4A, and CTCF. Binding affinities (SVM weights) of 10-mers were acquired from https://github.com/ljljolinq1010/A-robust-method-for-detecting-positive-selection-on-regulatory-sequences/blob/master/data/human_SVM_model/CEBPA/kmer_10_library_weigths.txt, https://github.com/ljljolinq1010/A-robust-method-for-detecting-positive-selection-on-regulatory-sequences/blob/master/data/human_SVM_model/HNF4A/kmer_10_library_weigths.txt, and https://github.com/ljljolinq1010/A-robust-method-for-detecting-positive-selection-on-regulatory-sequences/blob/master/data/human_CTCF_adaptation/human_SVM_model/all_merged_ctcf_kmer_10_library_weigths.txt for CEBPA, HNF4A, and CTCF, respectively. CEBPA and HNF4A are both liver-specific TFs; CTCF is expressed in multiple tissues and the binding sites studied are the union of the binding sites identified from multiple individual tissues (Liu and Robinson-Rechavi 2020). For each ancestral TFBS, we generated 1,000 random mutants as the control set. Each mutant was a two-step neighbor of the ancestral sequence and was generated as in the generation of the derived sequences. We simulated only 1,000 random mutants as the control set for each TFBS because the total number of two-step neighbors of a 10-mer is only 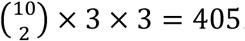. Positive selection is inferred for a TFBS if fewer than 10 sequences in the control set has higher SVM scores than that of the derived sequence (i.e., *P* < 0.01). Because the simulated TFBSs were subject to no selection, all positive cases identified in this analysis were false positives.

To mimic the ascertainment bias, we ranked the simulated TFBSs based on the SVM score of the derived sequence and removed the bottom 10%, 20%, … and 90% of TFBSs, respectively, obtaining nine subsets subject to increasing ascertainment biases. We then counted the number of TFBSs that showed *P* < 0.01 in the selection test in each subset to investigate the relationship between the ascertainment bias and the false positive rate.

### Reanalysis of human TFBSs

We examined the binding sites of human CEBPA, HNF4A, and CTCF, respectively. These TFBSs were previously studied by Liu and Robinson-Rechavi (2020), who used human (*Homo sapiens*) as the focal species and used human, chimpanzee (*Pan troglodytes*), and gorilla (*Gorilla gorilla*) to infer the sequence of the human-chimpanzee common ancestor. Data files analyzed here were made available by Liu and Robinson-Rechavi at https://github.com/ljljolinq1010/A-robust-method-for-detecting-positive-selection-on-regulatory-sequences/tree/master/data/human_deltaSVM (CEBPA and HNF4A binding sites) and https://github.com/ljljolinq1010/A-robust-method-for-detecting-positive-selection-on-regulatory-sequences/blob/master/data/human_CTCF_adaptation/human_deltaSVM/ctcf_deltaSVM_highertailTest.txt (CTCF binding sites).

We performed multiple testing corrections following the Benjamini-Hochberg procedure (Benjamini and Hochberg 1995) and computed adjusted *P*-values from *P*-values of the right-tail test in Liu and Robinson-Rechavi (2020). For the TFBS with the *i*^th^ smallest *P*-value in a given dataset, the adjusted *P*-value is 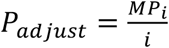, where *P*_*i*_ is the *P*-value reported by Liu and Robinson-Rechavi (2020) and *M* is the total number of tests to correct for.

We applied four methods to calculate the adjusted *P*-value, which we refer to as method 0, 1, 2, and 3, respectively. In method 0, *M* is simply the number of TFBSs in the dataset. In method 1, *M* equals twice the number of TFBSs with positive deltaSVMs in the dataset. Method 2 makes use of the relationship between the fraction of TFBSs removed (*F*_removed_) and the fraction of TFBSs with positive deltaSVMs (*F*_+_) in the remaining data, inferred from our simulated dataset by the regression *F*_removed_ = *kF*_+_ + *b. M* is then calculated by 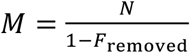, where *N* is the total number of TFBSs in the dataset. Values of *k* and *b*, along with inferred values of *F*_removed_ and *M* for the binding sites of the three human TFs are summarized in Table S3. In method 3, 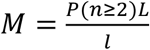, where *L* is the genome size, *l* is the mean length of TFBS sequences in the dataset, and *P*(*n* ≥ 2) is the probability that there are at least two substitutions between the ancestral TFBS and its derived version. *L* was set to be 6 × 10^9^, which is a rough estimate of human’s haploid genome size multiplied by 2 because a TFBS can be on either the Watson or Crick strand and *l* was computed from the empirical data to equal 269, 340, and 537 for CEBPA, HNF4A, and CTCF, respectively. *P*(*n* ≥ 2) was calculated as the ratio of the number of TFBSs used in the selection test (first row of Table 2) and the total number of TFBSs identified by ChIP-seq (16,212 for CEBPA, 27,782 for HNF4A, and 118,970 for CTCF).

Presumably, the missing TFBSs are unlikely to have small *P*-values. Thus, we assume that the ranks of *P*-values of TFBSs with *P* < 0.01 or *P* < 0.05 do not change upon the inclusion of missing TFBSs.

All simulations and analyses were conducted in an R environment (R Core Development Team 2010).

## ACKNOWLEDGEMENTS

We are grateful to S. Song and H. Xu for valuable comments. This work was supported by the research grant R35GM139484 from the U.S. National Institutes of Health to J.Z. D.J. was supported by the Rackham Predoctoral Fellowship of University of Michigan when performing this study.

---

**Figure S1.**
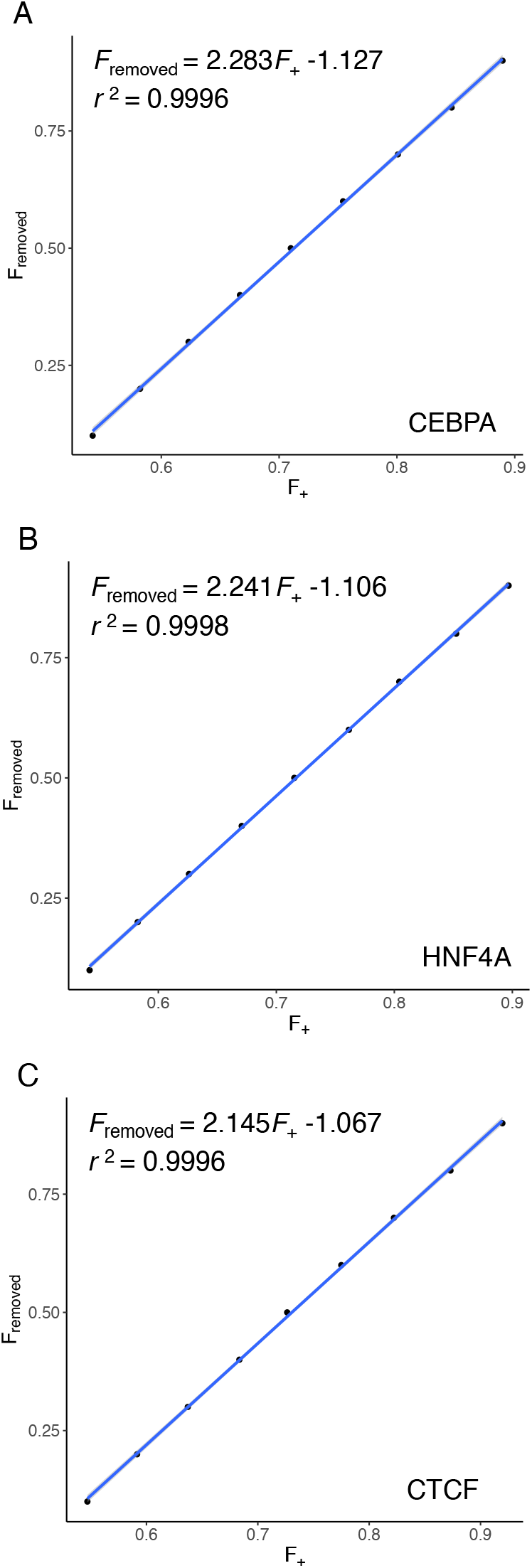
Linear relationship between the fraction of TFBSs with positive deltaSVMs (*F*_+_) and the fraction of TFBSs removed (*F*_removed_) in simulated data, based on binding affinity to human CEBPA (**A**), human HNF4A (**B**), and human CTCF (**C**), respectively.

**Table S1.** All simulated TFBSs and subsets with various fractions of the TFBSs with the highest derived SVM scores for human HNF4A. *P*-value in the last column is from a test of the null hypothesis that the false positive rate in a subset equals that in the complete dataset (two-sample *Z*-test for proportions).

| Subset | No. of TFBSs | No. of TFBSs with positive deltaSVMs | No. of false positives | *P*-value |
| --- | --- | --- | --- | --- |
| Complete dataset | 50,000 | 24,901 (49.80%) | 535 (1.107%) | N.A. |
| Top 90% | 45,000 | 24,378 (54.17%) | 535 (1.189%) | $4.4 \times 10^{-2}$ |
| Top 80% | 40,000 | 23,300 (58.25%) | 535 (1.338%) | $1.3 \times 10^{-4}$ |
| Top 70% | 35,000 | 21,909 (62.31%) | 535 (1.529%) | $2.2 \times 10^{-9}$ |
| Top 60% | 30,000 | 20,120 (67.07%) | 535 (1.783%) | $1.2 \times 10^{-17}$ |
| Top 50% | 25,000 | 17,881 (71.52%) | 535 (2.140%) | $1.7 \times 10^{-31}$ |
| Top 40% | 20,000 | 15,229 (76.15%) | 535 (2.675%) | $3.5 \times 10^{-55}$ |
| Top 30% | 15,000 | 12,063 (80.42%) | 535 (3.567%) | $1.4 \times 10^{-98}$ |
| Top 20% | 10,000 | 8,524 (85.24%) | 535 (5.350%) | $2.6 \times 10^{-191}$ |
| Top 10% | 5,000 | 4,484 (89.68%) | 535 (10.700%) | $< 10^{-300}$ |

**Table S2.** All simulated TFBSs and subsets with various fractions of the TFBSs with the highest derived SVM scores for human CTCF. *P*-value in the last column is from a test of the null hypothesis that the false positive rate in a subset equals that in the complete dataset (two-sample *Z*-test for proportions).

| Subset | No. of TFBSs | No. of TFBSs with positive deltaSVMs | No. of false positives | *P*-value |
| --- | --- | --- | --- | --- |
| Complete dataset | 50,000 | 25,024 (50.05%) | 548 (1.096%) | N.A. |
| Top 90% | 45,000 | 24,613 (54.70%) | 548 (1.218%) | $4.2 \times 10^{-2}$ |
| Top 80% | 40,000 | 23,663 (59.15%) | 548 (1.370%) | $1.1 \times 10^{-4}$ |
| Top 70% | 35,000 | 22,299 (63.71%) | 548 (1.566%) | $1.4 \times 10^{-9}$ |
| Top 60% | 30,000 | 20,504 (68.35%) | 548 (1.827%) | $4.9 \times 10^{-18}$ |
| Top 50% | 25,000 | 18,159 (72.64%) | 548 (2.192%) | $3.2 \times 10^{-32}$ |
| Top 40% | 20,000 | 15,499 (77.50%) | 548 (2.740%) | $1.7 \times 10^{-56}$ |
| Top 30% | 15,000 | 12,333 (82.22%) | 548 (3.653%) | $5.7 \times 10^{-101}$ |
| Top 20% | 10,000 | 8,730 (87.30%) | 548 (5.480%) | $5.2 \times 10^{-196}$ |
| Top 10% | 5,000 | 4,598 (91.96%) | 548 (10.960%) | $< 10^{-300}$ |

**Table S3.** Linear regression of the fraction of TFBSs removed (*F*_removed_) on the fraction of TFBSs with positive deltaSVMs in the dataset (*F*_+_).

| TF | Regression slope ($k$) | Intercept ($b$) | Inferred<br>$F_{\text{removed}}$ | Inferred total<br>number of TFBSs |
| --- | --- | --- | --- | --- |
| CEBPA | 2.283 | -1.127 | 29.19% | 8,200 |
| HNF4A | 2.241 | -1.106 | 20.30% | 15,977 |
| CTCF | 2.145 | -1.067 | 11.47% | 90,458 |

## Notes

### Competing Interest Statement

The authors have declared no competing interest.

